# Diabetes impacts endothelial Weibel-Palade body biogenesis and VWF secretion

**DOI:** 10.64898/2026.05.14.724893

**Authors:** Harriet J. Todd, Melissa Rose, Karen Forbes, Thomas A J McKinnon, Ramzi Ajjan, Marc A. Bailey, Lynn McKeown

## Abstract

Diabetes is associated with endothelial dysfunction, impaired wound healing, and increased thrombotic risk, yet the impact of diabetes on endothelial secretory organelles remains poorly understood. Weibel-Palade bodies (WPBs) are specialised endothelial granules that store and release von Willebrand factor (VWF) and other vasoactive cargo essential for haemostasis, inflammation, and vascular repair. Here, we investigated how diabetic environments influence WPB biogenesis and VWF structure under physiologically relevant flow conditions. Acute exposure of endothelial cells to constant or fluctuating high glucose concentrations, designed to model diabetic glycaemic conditions, did not alter WPB number or morphology under either static or high laminar shear stress conditions. In contrast, primary endothelial cells derived from a diabetic donor exhibited reduced Akt and eNOS signalling, significantly fewer WPBs, reduced intracellular VWF content, and shorter stimulus-evoked VWF strings compared with non-diabetic endothelial cells. Although total cellular VWF levels were reduced, high molecular weight (HMW) VWF content within endothelial lysates was not significantly altered. Plasma from diabetic patients demonstrated elevated circulating VWF levels together with marked inter-patient heterogeneity in VWF multimer composition. These findings suggest that chronic diabetes-associated endothelial dysfunction, rather than hyperglycaemia alone, alters WPB biology and VWF handling. We propose that dysregulated basal endothelial secretion may deplete endothelial VWF stores, limiting appropriate stimulus-coupled WPB release during vascular injury and contributing to defective vascular repair in diabetes.

## Introduction

Diabetes Mellitus (DM) represents a major and growing global health burden, affecting over 500 million people worldwide and conferring a substantially increased risk of cardiovascular disease, microvascular complications, and impaired tissue repair ^2^. Among the most debilitating clinical consequences of diabetes is defective wound healing, which contributes to chronic ulceration, infection, and lower-limb amputation ^3,4,5^. Although hyperglycaemia, inflammation, and metabolic dysregulation are well-established drivers of diabetic vascular pathology, the cellular mechanisms linking the diabetic vascular environment to impaired haemostatic and reparative responses remain incompletely understood.

The vascular endothelium plays a central role in coordinating haemostasis, inflammation, and angiogenesis during tissue repair ^6 7^. A key feature of endothelial cells is the presence of Weibel-Palade bodies (WPBs), specialised secretory organelles that store and release a diverse array of vasoactive and haemostatic cargo ^8 9^. The most abundant component of WPBs is von Willebrand factor (VWF), a multimeric glycoprotein essential for platelet adhesion and clot formation at sites of vascular injury ^10^. In addition to VWF, WPBs package inflammatory mediators and angiogenic factors including P-selectin, angiopoietin-2 (Angpt2), and interleukin-8, enabling rapid, stimulus-dependent secretion in response to vascular stress or injury. Through regulated exocytosis of this cargo, WPBs play a critical role in initiating haemostasis, recruiting immune cells, and supporting vascular remodelling during wound healing. Importantly, WPBs are not a homogeneous population ^11 12^; their biogenesis, morphology, and functional output are highly sensitive to the endothelial microenvironment ^11 13^. WPB length and cargo organisation are determined during biogenesis at the trans-Golgi network and are influenced by intracellular signalling pathways and external cues ^14 15 16^. We and others have previously demonstrated that haemodynamic forces, particularly shear stress generated by blood flow, are key regulators of WPB formation and function ^15^. Under physiological laminar flow, endothelial cells exhibit altered WPB morphology and reduced storage of ultra-large VWF multimers, consistent with a more quiescent, antithrombotic phenotype. In contrast, disturbed or low shear conditions promote the formation of elongated WPBs enriched in pro-thrombotic and pro-inflammatory cargo.

Diabetes is associated with profound alterations in vascular haemodynamics, including endothelial dysfunction, reduced vessel compliance, and disturbed flow patterns, particularly within the microvasculature ^17 18^. These changes are exacerbated in tissues prone to impaired healing, such as the lower extremities, where abnormal shear stress and perfusion are hallmarks of diabetic vascular disease. Despite this, the impact of diabetes-associated flow disturbances on WPB biogenesis and secretory function has not been defined although evidence from some animal models suggest changes.

Here, we hypothesised that diabetes-relevant haemodynamic environments alter WPB formation and function, thereby contributing to defective endothelial responses during wound healing. By integrating in vitro flow models with physiologically relevant endothelial cells or endothelial cells exposed to culture conditions that mimic the diabetic glycaemic environment, this study investigates how diabetes influences WPB morphology and secretory capacity. Defining how diabetes-associated vascular environments reshape WPB biology will provide new mechanistic insight into endothelial dysfunction and identify potential targets to improve vascular repair in diabetes. ^19^

## Methods

### Cell culture

HUVECs: Pooled Human Umbilical Vein Cells (HUVECs) (PromoCell) cultured in basal endothelial cell growth media with supplement pack (PromoCell) and antimycotic (Gibco) added forming endothelial growth medium (EGM-2).

hAECS: Primary human aortic endothelial cells (hAECs: lot.483z027.6) or dhAECs (lot.4727002.1) from a patient formally diagnosed with type II diabetes (PromoCell) were cultured in Endothelial Cell Growth Medium MV2 (EMV-2) supplemented with Growth Medium MV2 supplement pack (PromoCell) and antimycotic (Gibco).

All cells were maintained in a 5% CO_2_ environment in a 37°C incubator and utilised between passages 0-5.

### Shear stress

All experimental cells were subjected to laminar high sear stress (HSS) for the timeframe of the experiments unless otherwise stated. 100 ml of cells at 7.5 x10^5^ cells/ml plated on µ-Slide I 0.4 Luer (Ibiditreat) (IBIDI) and incubated for 18-24 hrs prior to flow. The IBIDI pump system generated unidirectional laminar flow at 5 dyn/cm^2^ for the first hour, increased to 10 dyn/cm^2^ for 48 hrs. Each condition had a static counterpart incubated for 48 hrs at 37°C 5% CO_2_ with media changed at 24 hrs.

### Diabetic models

#### High glucose model

Standard EGM-2 (glucose [5.5 mM]) was supplemented with 19.5 mM of filtered D-glucose solution ((Merck), final concentration of 25 mM. Molarity confirmed using GlucCell® glucose monitoring system (Merck). 19.5 mM mannitol (D-mannitol (M4125, Sigma-Aldrich, UK)) was used as an osmolarity control. Cells were cultured in a constant glucose concentration for 48 hours.

#### Ambulatory glucose model

Refer to Fig. 1. Model based on data from continued glucose monitoring, replicating the extremes in glucose fluctuations throughout an ‘average’ 8-hour day with ‘average’ eating habits ^1^. Custom glucose free EGM-2 media (Promocell) was adapted to reflect extreme clinically relevant hypo and hyperglycaemic environments in either diabetic or control conditions. EMG-2 was supplemented with filtered D-glucose (Merck) to stated concentrations. The diabetic model was comprised of low glucose at 2.5 mM and high at 20 mM conditions compared to normal control of low glucose at 5.5 mM and high at 10 mM.

**Figure 1.**
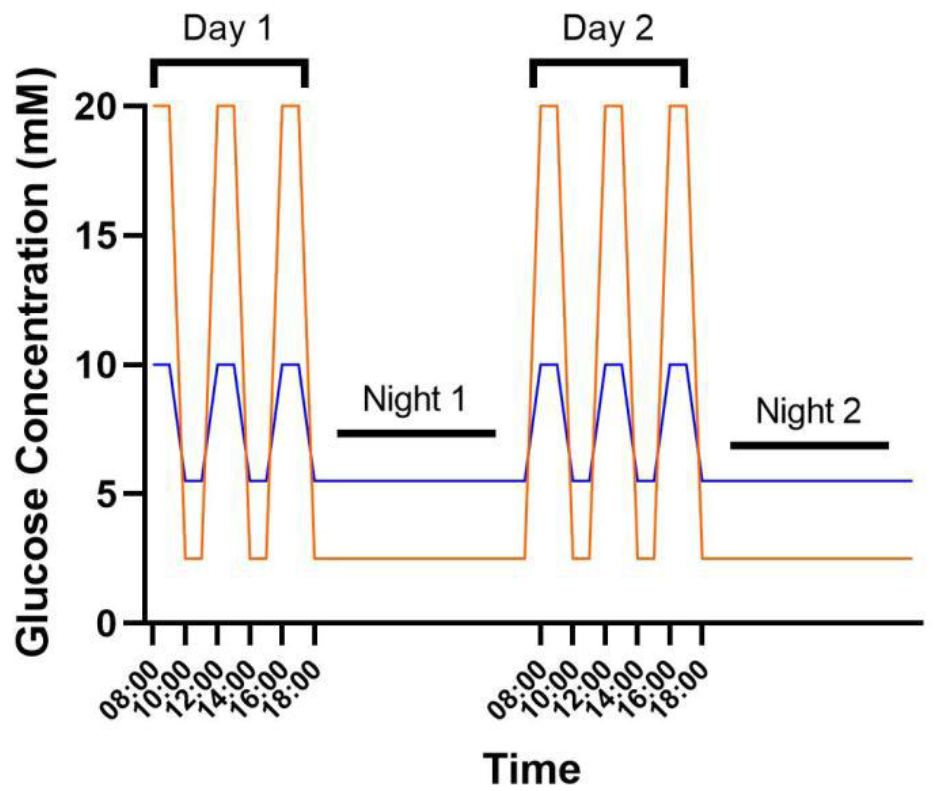
Fluctuating glucose modelling diabetic glycaemic levels or normo-glycaemic levels in a 48-hour cycle. Whilst under HSS for 48 hrs, endothelial cells were exposed to glucose cycles of hypo and hyperglycaemic environment. Day 1 and 2 consisted of two ten-hour periods to mimic an ‘average’ day of blood glucose fluctuations and an overnight hypoglycaemic cycle. Concentrations of glucose were extrapolated from continuous glucose monitoring data from Battelino et al. ^1^ to maintain physiological relevance. Control hyperglycaemia (post prandial) was set at 10 mM, hypoglycaemia was 5.5 mM, diabetic hyperglycaemia was 20 mM and hypoglycaemia 2.5 mM.

During 48 hrs of HSS cell media fluctuated through 3 x cycles. The first and second cycles consisted of 2 hrs of the relevant high glucose levels switching to 2 hrs of the corresponding low glucose condition, the third cycle consisted of the high glucose level for 2 hrs and the relevant low glucose overnight to reflect the time in which a person would typically be asleep.

### VWF string formation

After 48 hrs of HSS, slides were subjected to 2.5 dyn/cm^2^ laminar flow 10 mins and stimulated with 150 µM of histamine (Sigma) or 1 µM PMA (Promega) allowing VWF secretion and string formation. Slides then fixed (4% PFA) before immunofluorescent staining as per below.

### Immunofluorescent staining

At 48hrs, cells were fixed (4% PFA 10 minutes), permeabilised 10 minutes (0.1% triton) and incubated with monoclonal anti-VWF primary antibody (Dako) for 1 hr. Detection was performed by incubation with anti-mouse secondary antibody (Alexa Fluor 594 anti-mouse: Jackson ImmunoResearch Labs) 30 minutes. Nuclei were stained with Hoechst 33342 for 7 minutes, and slides mounted using Prolong gold mounting media (Invitrogen). For VWF strings detection, the permeabilization step was omitted as they are expressed extracellularly. Platelets were stained with CD41 primary antibody and corresponding secondary after fixation.

### Microscopy

An Olympus IX83 widefield fluorescent microscope and x60 oil objective was used to image 10x independent randomised fields of view (FOV). Olympus CellSens software was used to set the acquisition parameters where the optimum exposure was established for DAPI (blue) and RFP (red) fluorescence filters on controls then applied equally to all slides per biological repeat. z-stacks acquisition per FOV composed of 10x 0.2 µm steps. Each z-stack underwent constrained iterative (5x) deconvolution using a proprietary algorithm to produce a maximum intensity projected image.

A confocal LSM880 Inverted Zeiss LSM880 microscope was used to visualize VWF strings in 5x randomised FOVs using a x4[air objective. Zeiss ZEN black software, controlled acquisition and the optimum gain was applied to each image taken on all slides per biological repeat. Z-stack images were acquired for each FOV, 5x focal planes taken at 0.2 µm steps. Images were tiled (2×2) ensuring longer VWF strings were captured. Maximum intensity projections were generated and a medium stitch applied ensuring continuity between tiles.

### Image analysis

FIJI (Fiji is just ImageJ) was used to analyse maximum intensity projections. A macro was employed for analyses to prevent bias: macro 1 (Appendix I supplementary) enabled equal segmentation of WPBs and measurement of WPB number and feret size (length). Number of nuclei were used to calculate the average number of WPB per cell in each FOV (total WPBs divided by number of nuclei). To analyse the VWF strings macro 2 (Appendix 2 supplementary) was utilised, removing background ‘noise’ fluorescence and assessing number and length of VWF strings in each FOV.

### Western blots

#### hAEC and dhAEC cell lysis

Cells lysed with 100 µl cold lysis buffer (NP-40 Thermo scientific) containing 1:500 µl anti-protease cocktail (Sigma-Aldrich, P8340), and 1:50 µl broad-range anti-phosphatase (Sigma-Aldrich, 524629). Lysate collected, spun 10 min (12,000 x g) and supernatant collected and frozen.

#### Blot

Following BCA protein quantification (Rapid Gold BCA Kit A53225, ThermoFisher Scientific), samples were prepared to equivalent protein concentrations in sample buffer (200 mM Tris (pH 6.8), 8% SDS, 40% glycerol, 8% beta-mercaptoethanol, and 0.1% bromophenol blue), heated at 95°C 5 mins. Samples loaded onto precast 4-20% acrylamide gels (Bio Rad mini-PROTEAN TGX) with 4 µl of molecular weight protein ladder (Precision Plus Protein Standard Dual colour Bio-Rad, #161-0374). Gels run at 160V for ∼90 mins in 1x TGS running buffer (diluted from 10x with dH_2_O, Bio-Rad). Transfer was performed using the HMW programme on a BioRad turbo system for 20 mins, after which PDVF membranes were blocked using 5% milk PBS-Tween 1 hr. After washing, membranes were incubated with primary antibodies (BD Biosciences: 1:1000 mouse anti-pAkt. Cell Signalling: 1:1000 rabbit anti-pAkt, 1:1000 mouse anti-teNOS or 1:500 mouse anti-peNOS) in PBS-T overnight. Following 3x 10 mins washes (1xT-BST), membranes incubated 30 mins with secondary antibodies (Jackson Immunoresearch: goat anti-mouse or anti-rabbit HRP). Detection of signal was obtained using SuperSignal West Dura Extended Duration Substrate ECL (ThermoFisher Scientific) and imaged using an iBright FL1500 (ThermoFisher, Scientific, UK).

### VWF multimer gels

Cells were cultured as stated for 48 hours prior to lysis with RIPA buffer (Pierce Thermo Scientific). Protein content was quantified using Rapid Gold BCA Kit (A53225, ThermoFisher Scientific) as per manufacturer’s instructions and lysate protein concentration equalized in sample buffer (10 mM Tris, 10 mM EDTA, 2% (w/v) SDS, 0.03% (w/v) bromophenol blue, 15% glycerin pH 8.0). The reference plasma (normal pooled plasma [First Link UK, 2800850]) and plasma samples were diluted 1:10. Samples were denatured at 60°C for 30 mins and loaded, 10 µl/ well, into Tris gels (200 mM Tris, 100 mM Glycine, 0.1% SDS, pH 9.0: 1.2 % high gelling temperature agarose (Seakem Lonza, USA). Gels were run at 4°C in running buffer (100 mM Glycine, 100 mM Tris and 0.1% SDS pH 8.4) at 65 V for 20 mins then 25V for 3-5 hrs. Gels were left in DDT (Dithiothreitol: BioRad, UK) for 20 mins prior to transfer. Transfer was performed using the HMW programme on a BioRad turbo system for 20 mins, after which membranes were blocked using 5% milk PBS-T 1 hr. After washing with PBS-Tween, membranes were incubated 1:5000 rabbit anti-human VWF-HRP (Dako) in PBS-T overnight. Detection of signal was obtained using SuperSignal West Dura Extended Duration Substrate ECL (ThermoFisher Scientific) and imaged using an iBright FL1500 (ThermoFisher, Scientific, UK). Auto-exposure determined the optimum imaging time, but increasingly higher exposures ensured all bands were captured.

### Analysis of multimer gels

Gels were analysed using ImageJ/ FIJI. Briefly, images were inverted and the line tool used to measure the intensity of each VWF band down the column. The intensity profile plot was generated by highlighting all the lines and using the multi plot feature in the ROI manager. The ImageJ generated list of values were used to calculate the area under the curve (AUC) for each profile by using a trapezoidal integration. The AUC of each profile was normalized to the reference plasma for each gel. High molecular weight AUC determined after the position of the 5^th^ LMW peak.

### Statistics

All microscopy images were acquired in a random manner and analysis equally automated using Fiji macros and therefore no blinding was necessary. All averaged data are presented as mean ± SEM. One-way or two-way ANOVA was performed as appropriate to determine whether significant differences existed among three or more groups, coupled with Bonferroni post hoc test. Two sample t-test was performed when comparing two sample data groups after normality was approached by a log-transformation, and equal variance confirmed. Significance values are shown in the figures or in figure legends. Statistical significance was considered to exist at probability *p* < 0.05, (*** < 0.001, ** < 0.01, * < 0.05). Where comparisons lacked an asterisk or marked NS, no significant difference between groups was observed. OriginPro 2020 software and R studio was used for data analysis and presentation. *n / N* = number of independent repeats / numbers of technical repeats.

## Results

### High glucose alone is insufficient to alter WPB number or morphology in endothelial cells

To determine whether hyperglycaemia per se is sufficient to influence WPB biogenesis, human umbilical vein endothelial cells (HUVECs) were cultured for 48 hrs in media containing high glucose (20 mM) and compared to cells maintained under normo-glycaemic conditions (5 mM: control). Cells were subjected to either static conditions or high laminar shear stress (HSS) for 48 hours to ensure that potential glucose-dependent effects were not masked by static culture artefacts. Following shear exposure, endothelial cells were immunostained for VWF to visualise WPBs, and WPB number and length were quantified using high-resolution microscopy (Fig. 2A-C). Representative images (Fig. 2A) and quantitative analysis revealed no significant effect of high versus normo-glycaemia in the number of WPBs per cell (Fig. 2B) nor the length of WPBs (Fig 2C). Different constant glucose concentrations had no detectable effect on the endothelial response to high laminar shear stress (HSS), as cells exposed to either normo-glycaemic or high-glucose conditions showed comparable reductions in WPB length following shear exposure.

**Figure 2.**
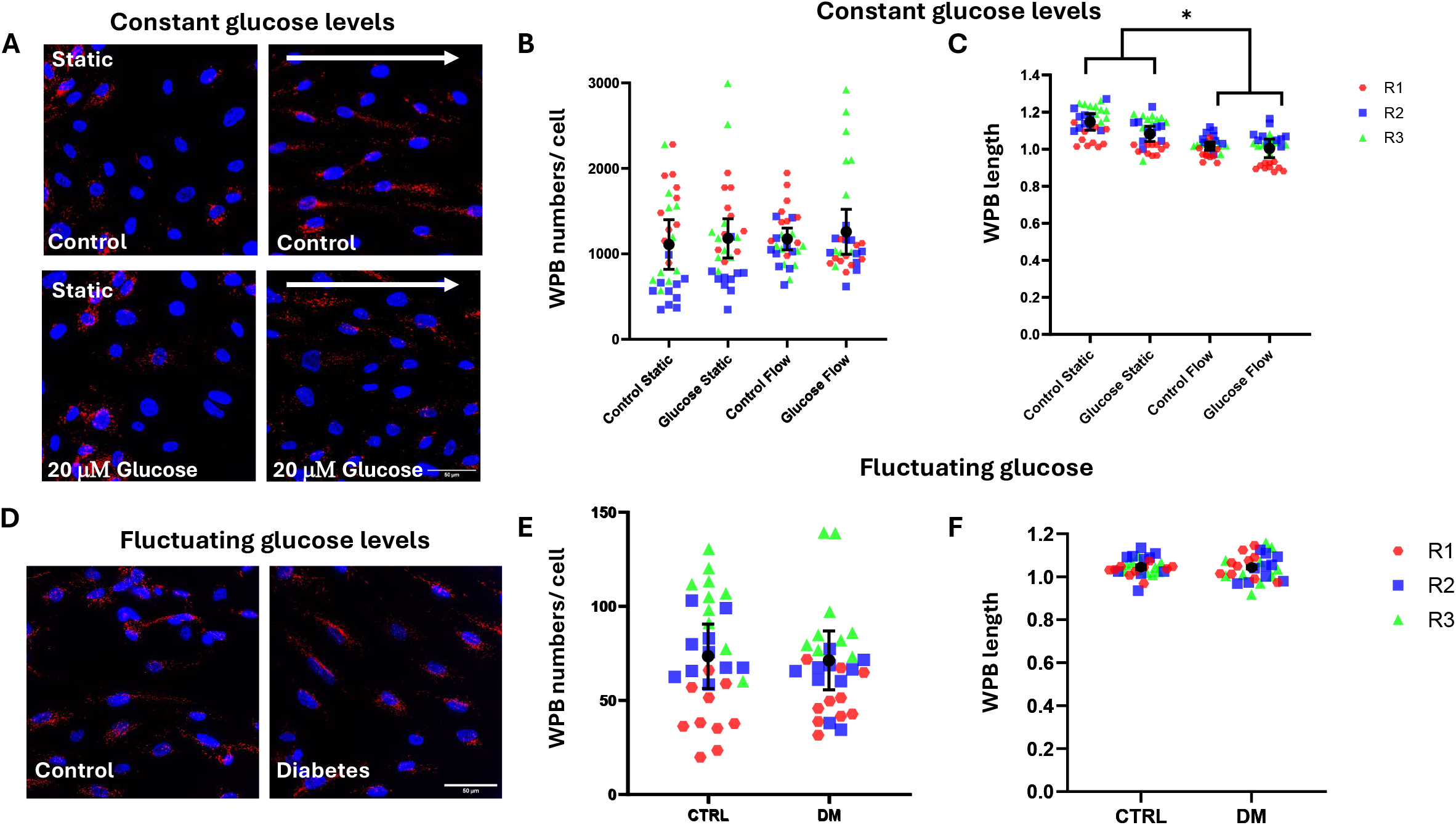
High glucose exposure does not alter WPB number or morphology in endothelial cells. (A) Representative immunofluorescent images of VWF (red) in Human Umbilical Vein Endothelial Cells (HUVECs) cultured for 48 hours under normo-glycaemic conditions (5 mM: control) or constant high glucose (20 mM). Cells were exposed to defined high laminar shear stress (HSS: arrow) or cultured under static conditions for 48 hours. The control condition includes 15 mM mannitol to equalize viscosity and osmotic effects. (B, C) Mean data quantified showing the number of WPBs per cell and WPB (feret) length. (D) Representative images of VWF (red) in HUVECs cultured under a 48 cycle of fluctuating glucose levels designed to mimic fluctuations observed in diabetes (DM) or normo-glycaemic (CTRL) conditions. (E-F) Quantification of WPBs per cell and WPB length. No significant differences were observed in numbers or length across glucose treatments. Data represent mean +/- SEM, n / N = 3 / 30 experiments; statistical analysis performed using two-way ANOVA. Scale bar = 50 μm.

To better reflect physiological exposure patterns, HUVECs cultured under HSS were subjected to fluctuating glucose conditions designed to mimic glycaemic variation over a 48-hour cycle as observed in either diabetic or non-diabetic individuals (regimen shown in Fig. 1 taken from constant glucose monitoring). Consistent with the constant glucose conditions, no differences were observed in either WPB numbers or WPB length in cells exposed to fluctuating high glucose compared to normo-glucose containing media (Fig. 2D-F) Osmotic controls (mannitol) were included to exclude confounding effects of altered osmolarity and showed no detectable impact.

Collectively, these data indicate that acute exposure to diabetic-range glucose concentrations, even under physiologically relevant shear stress, is insufficient to alter WPB biogenesis or morphology in endothelial cells.

### Endothelial cells from diabetic patients exhibit altered signalling and reduced WPB abundance

Given the absence of a direct hyperglycaemic effect, we next examined whether endothelial cells derived from diabetic patients exhibit intrinsic differences in WPB biology. Human aortic endothelial cells (hAECs: physiological relevant endothelial cells that also change their WPB biogenesis in respond to flow ^15^) isolated from a diabetic donor (dhAECs) displayed significant alterations in key metabolic and endothelial signalling pathways compared to non-diabetic controls (Fig. 3A, B). Immunoblot analysis demonstrated reduced phosphorylation of Akt (pAkt) together with reduced endothelial nitric oxide synthase (eNOS) levels, consistent with impaired insulin signalling and nitric oxide bioavailability characteristic of diabetic endothelium ^18^.

**Figure 3.**
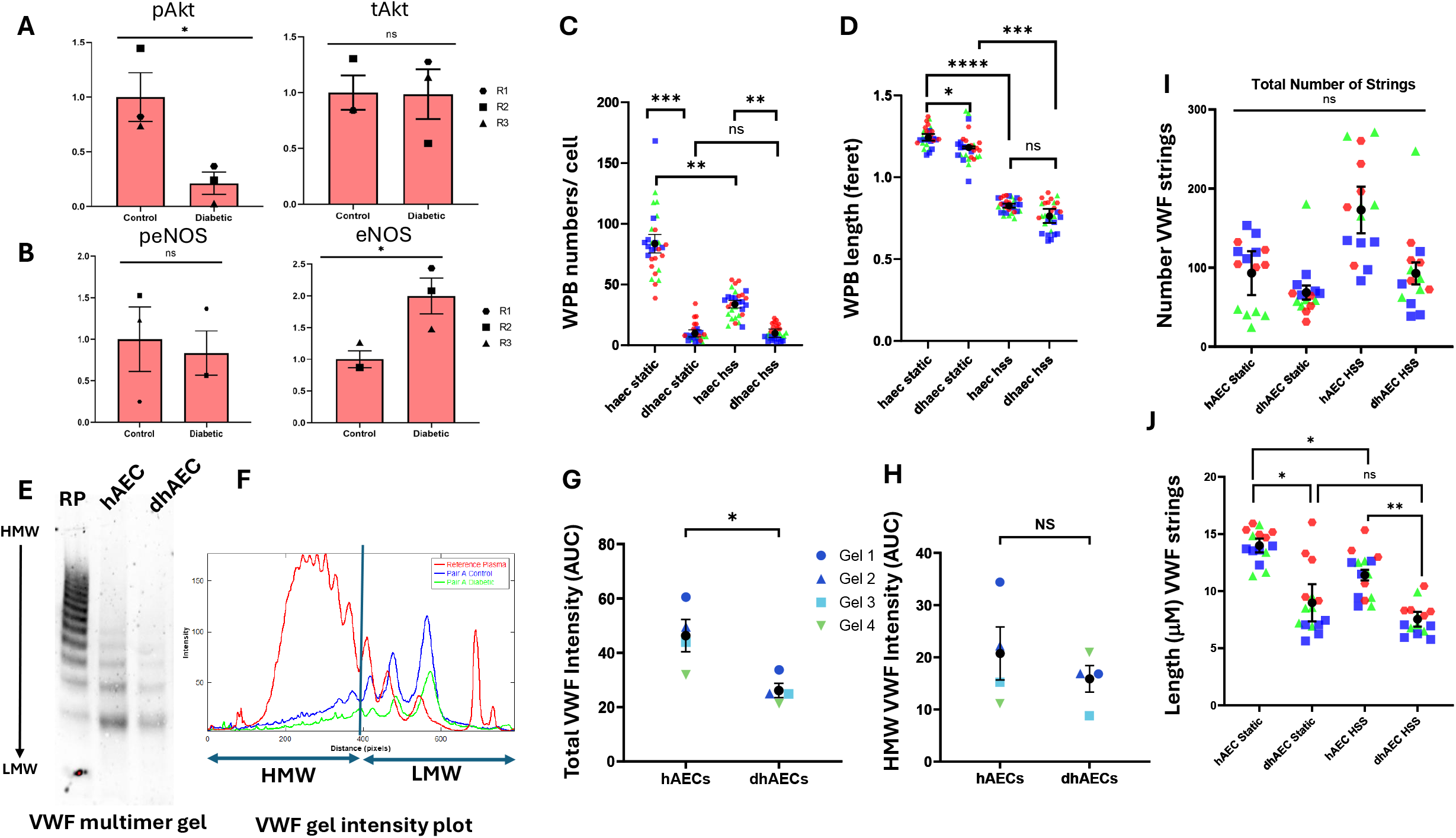
Endothelial cells from diabetic donors exhibit altered signalling, reduced WPB abundance, and altered VWF multimer composition. (A) Mean densitometric analysis of Akt normalised to β-actin and phospho-Akt (pAkt) normalised to total Akt measured in lysates from hAECs (control) and dhAECs (diabetic). (B) Mean densitometric analysis of total eNOS and phospho-eNOS (peNOS) measured in lysates isolated from non-diabetic and diabetic hAECs. Diabetic hAECs (dhAECs) exhibit reduced pAkt and total eNOS relative to total protein levels. n = 3 for all gels. (C) Mean data demonstrating the number of WPBs per cell in control hAECs versus diabetic hAECs maintained under static or laminar high shear stress (HSS) conditions for 48 hrs. WPB numbers are significantly reduced in dhAECs under both static and HSS conditions compared to hAECs. (D) Mean data of the feret length of WPBs in hAECs and dhAECs maintained in static or HSS conditions for 48 hrs. All imaging data are represented as mean +/- SEM from n / N = 3 / 30. Statistical analysis performed using unpaired t-test. (E) Representative VWF multimer blot from non-diabetic and diabetic hAEC lysates. RP = reference plasma. (F) Representative intensity profile from the bands shown in gel (E), each peak represents the intensity of each band. Low molecular weight (LMW) VWF consists of bands 1 – 5. HMW is determined after the 5^th^ LMW band. (G) Mean data +/- SEM of the area under the curve (AUC) of the intensity profiles indicating that total VWF is reduced in lysates from dhAECs compared to hAECs. (H) Mean data of the AUC of HMW VWF in hAECs compared to dhAECs. Data represented as the mean +/- SEM from n = 4. Statistical analysis was performed using unpaired *t*-test where * = p < 0.05, ** = p < 0.01. (I) Mean data of the number of VWF strings and (J) VWF string length following histamine stimulation. Analysed using unpaired *t*-test * = *p* <0.05, ** = *p* <0.01. n / N = 3 / 15.

Quantitative analysis of WPBs using high-resolution immunofluorescence imaging of VWF demonstrated a marked reduction in WPB numbers per cell in diabetic hAECs compared with non-diabetic controls (Fig. 3C). This reduction was observed under both static and HSS conditions, indicating that physiological shear does not rescue the defect. Consistent with our previous observations ^15^, exposure to HSS induced the formation of shorter WPBs in both control and diabetic cells (Fig. 3D). Under static conditions however, WPB length was more variable and significantly reduced in diabetic cells.

Given that endothelial signalling pathways downstream of Akt and eNOS regulate WPB biogenesis and secretory capacity, and that both pathways were reduced in diabetic hAECs (Fig. 3A, B), we next asked whether the decrease in WPB number reflected altered total cellular VWF levels and/or changes in VWF multimer composition. Multimer gel analysis was performed using equivalent total protein loading (Fig. 3E). Since multimer profiles from cell lysates differ from plasma (see reference plasma control in Fig. 3E and F compared to lysates which lack several of the highest-order HMW multimers), peak ratio-based comparisons are less reliable. Instead, quantification of the area under the curve (AUC) was used to assess relative VWF content. Consistent with the morphological findings, biochemical analysis demonstrated reduced total cellular VWF levels in diabetic hAECs (Fig. 3E-G), indicating that the reduction in WPB abundance was accompanied by decreased intracellular VWF content rather than a global change in total protein abundance.

Because VWF function depends on both total levels and multimer size, with high molecular weight (HMW) multimers exhibiting greater platelet-binding capacity and haemostatic activity ^15 20^, we next assessed multimer distribution. Quantification of the area under the curve (AUC) to assess relative HMW content after the 5^th^ low molecular weight (LMW) peak demonstrated similar levels in control and diabetic hAECs (Fig. 3H).

To determine whether reduced total cellular VWF levels in the dhAECs translated into functional consequences, we examined VWF string formation following agonist stimulation. Although the number of strings formed was not significantly altered, diabetic hAECs produced significantly shorter VWF strings than non-diabetic controls, suggesting altered VWF packaging and/or secretion dynamics (Fig. 3I).

Together, these findings indicate that diabetes is associated with altered signalling, reduced VWF levels and decreased WPB abundance consistent with diminished haemostatic reserve.

### Circulating plasma VWF is elevated in diabetes but exhibits heterogeneous multimer composition

Interpretation of endothelial cell phenotypes derived from diabetic donors is inherently limited by inter-individual variability and incomplete clinical metadata, including differences in disease duration, glycaemic control, and treatment history. Such heterogeneity complicates attribution of observed cellular defects solely to diabetes and raises the possibility that disease severity or therapeutic exposure may differentially influence WPB biogenesis and function.

To determine whether our cellular observations were reflected at the population level, we analysed plasma VWF from a cohort of 21 patients with diabetes and compared them with non-diabetic controls (Fig. 4). Total circulating VWF levels were significantly elevated in the diabetic cohort, consistent with established evidence of endothelial activation or dysfunction in diabetes (Fig. 4A-C). Despite this overall increase, multimer analysis revealed substantial inter-individual variability in VWF multimer composition within the diabetic cohort. Both higher-molecular-weight (HMW) and lower-molecular-weight (LMW) species varied between individuals, but no consistent shift towards a uniform multimer distribution was observed across the cohort (Fig. 4D-F). Some patients exhibited relative depletion of HMW multimers, whereas others showed preservation or enrichment of these species.

**Figure 4.**
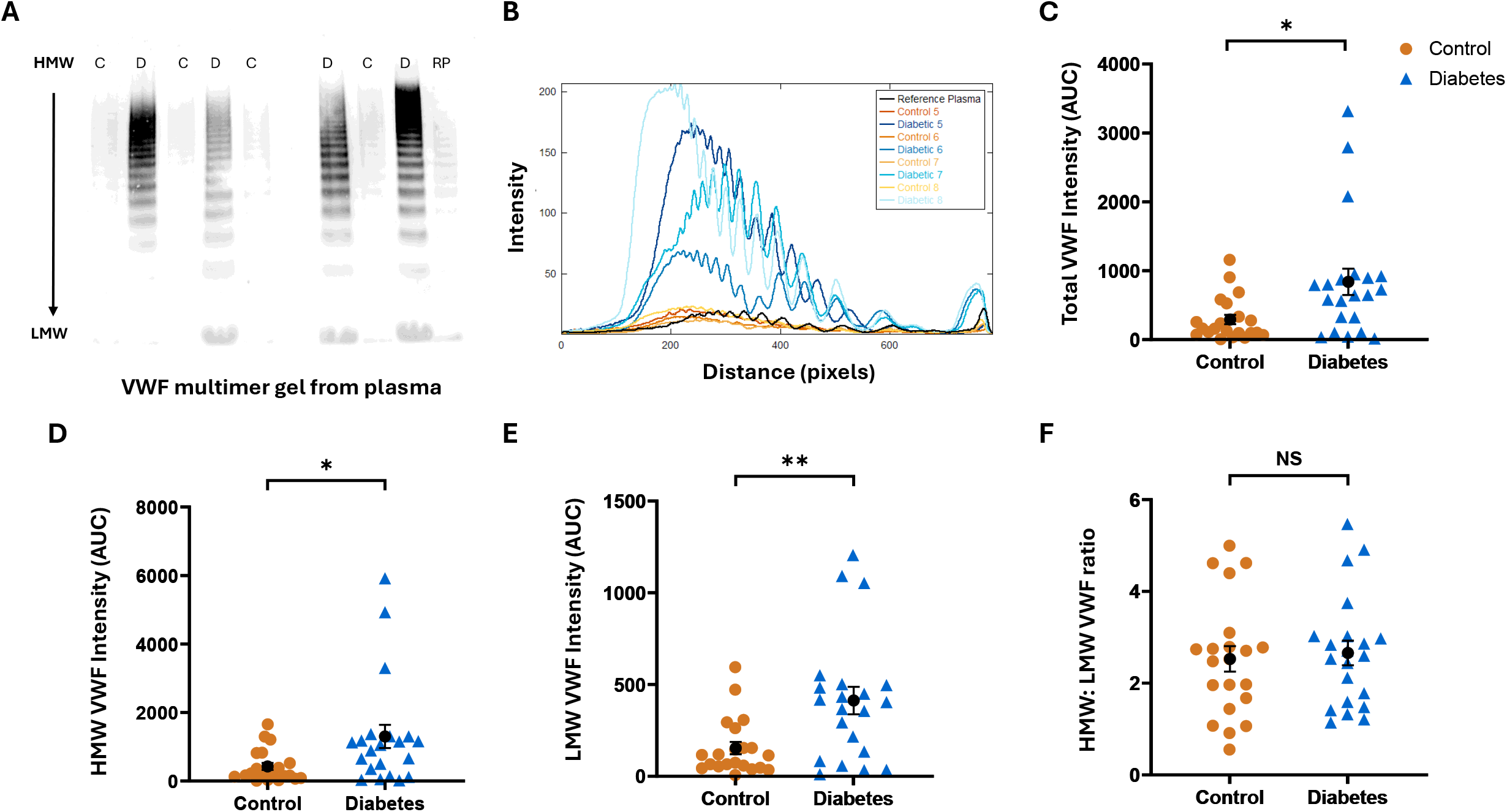
Circulating VWF is elevated in plasma from diabetic patients but exhibits heterogeneous multimer composition. (A) Representative VWF multimer blots of the plasma taken from control patients (C: non-diabetic) and patients with diabetes (D). RP = reference plasma. (B) Representative multimer band intensity profiles demonstrating variable multimer distribution across diabetic patients with reduced VWF levels. (C) Quantification (mean +/- SEM) of AUC of the intensity profiles representing total circulating VWF levels and (D) (mean +/- SEM) of AUC of the intensity profiles representing HMW circulating VWF levels and (E) LMW VWF levels. (F) HMW/LMW ratio (mean +/- SEM) of VWF multimers. Plasma VWF levels are significantly elevated in diabetes; however, multimer analysis reveals marked inter-patient variability with no uniform shift in multimer pattern. Analysed using unpaired *t*-test * = *p* <0.05, ** = *p* <0.01. n = 21/cohort.

The divergence between reduced intracellular VWF stores and elevated circulating plasma VWF suggests that diabetes alters basal endothelial secretion and/or clearance of VWF rather than simply reducing biosynthetic capacity. Together these results highlight a complex relationship between diabetes, WPB biology, and systemic VWF homeostasis.

## Discussion

In this study, we investigated how diabetic environments influence endothelial WPB biogenesis, cargo organisation, and VWF homeostasis. Our data demonstrate that acute exposure to constant high or fluctuating high/low glucose levels, as observed in diabetes, does not alter WPB number or morphology, whereas endothelial cells derived from a diabetic patient exhibit altered metabolic signalling, reduced WPB abundance, and changes in VWF content and structure. At the population level, plasma from diabetic patients contained elevated VWF levels, albeit with considerable heterogeneity in multimer organisation. Together, these findings suggest that diabetes-associated endothelial dysfunction, rather than hyperglycaemia alone, underlies dysregulated WPB biology and VWF handling.

A major strength of this study is the integration of both static and flow-based culture systems. Endothelial cells in vivo are continuously exposed to shear stress, which is a key determinant of endothelial phenotype and WPB formation. By comparing static and flow conditions, we were able to distinguish glucose-dependent effects from those influenced by haemodynamics and demonstrate that reduced WPB abundance in diabetic endothelial cells persists even under physiological shear stress. The use of parallel systems therefore enhances the physiological relevance of our findings.

Nevertheless, there are limitations associated with in vitro flow models. The use of Luer slides and the Ibidi flow system allows precise control of shear stress but does not fully recapitulate the complex and heterogeneous flow patterns present in diabetic vasculature, particularly in the microcirculation. In addition, long-term culture under flow is technically challenging and may limit the ability to model chronic disease states. Despite these constraints, flow-based approaches remain essential for studying WPB biology and represent a significant advance over static culture alone.

The absence of an effect of constant or fluctuating high glucose on WPB formation in our experimental model likely reflects the relatively short duration of glucose exposure. Endothelial cells were cultured under hyperglycaemic conditions for three days, which may be insufficient to induce the sustained metabolic and signalling defects characteristic of chronic diabetes. Consistent with this interpretation, endothelial cells derived from a patient with diabetes exhibited reduced Akt and eNOS phosphorylation together with fewer WPBs, supporting the notion that prolonged metabolic dysfunction, rather than acute glucose elevation alone, is required to alter WPB biogenesis. These observations suggest that changes in WPB number and cargo composition observed in diabetic endothelial cells, including the selective reduction in high-molecular-weight VWF, are more likely to reflect long-term disease-associated reprogramming than short-term glucose exposure per se. Future studies incorporating extended metabolic conditioning or additional diabetes-relevant stressors, such as oxidative stress or inflammatory signalling, will be important to resolve this discrepancy. Moreover, because endothelial turnover and repair are altered in diabetes, hyperglycaemia may exert part of its effect upstream at the level of endothelial progenitor cells within the bone marrow, where chronic metabolic stress has the potential to reprogramme endothelial lineage commitment and thereby influence the phenotype and secretory capacity of newly generated endothelial cells entering the circulation.

The use of primary endothelial cells from diabetic patients represents an important physiological strength of this work. However, reliance on cells from single donors introduces variability and limits statistical power, particularly in the absence of detailed clinical metadata such as disease duration, glycaemic control, or medication history. Moreover, the limited availability of patient-derived endothelial cells makes it challenging to obtain the biological numbers required for robustly powered mechanistic studies. In contrast, plasma samples are more readily available and can be collected alongside relevant clinical information, allowing increased experimental power and stratification.

Our plasma analysis revealed consistently elevated VWF levels in diabetic patients, in line with a prothrombotic vascular phenotype. This increase may help explain the heightened thrombotic risk associated with diabetes. However, paradoxically, diabetes is also characterised by poor wound healing. One possible explanation is impaired WPB trafficking leads to constitutive or dysregulated VWF release. Continuous basal secretion of VWF could deplete endothelial stores, such that when acute, stimulus-driven secretion is required during vascular injury or wound repair, the characteristic bolus release of WPB cargo is blunted or absent.

In future work, it will be essential to stratify diabetic samples based on disease duration, glycaemic control, and therapeutic regimens to better understand how these variables influence WPB biology and VWF homeostasis. Such studies will be critical for defining how chronic diabetes reshapes endothelial secretory function and for identifying potential strategies to restore appropriate WPB-mediated responses in vascular disease and impaired wound healing.

## Supporting information

Supplemental macros

## Acknowledgements

This research was supported by a BHF PhD studentship and a Leeds 4yr PhD studentship (FS/4yPhD/F/21/34153 and FS/4yrPhD/24/34209) both as part of a 4yr BHF PhD programme awarded to Harriet Todd and Melissa Rose. We’d like to thank the University of Leeds, Faculty of Biological Sciences Bioimaging facility for their advice and use of facilities (https://biologicalsciences.leeds.ac.uk/facilities/doc/bio-imaging-flow-cytometry). We would like to thank Prof. David Beech for helpful discussions on the manuscript.

## Author Contributions

HJT performed the experiments. HJT, MR, KF, MB, and LM analysed the data and created the graphs. TM provided the training and resources for the multimer gel analysis. RA collected and provided the clinical samples. MR created the graphical abstract. LM conceived the study and co-wrote the article with input from all authors. LM generated research funds and supervised the PhD studentships.

