## Supplemental macros for "Diabetes impacts endothelial Weibel-Palade body biogenesis and VWF secretion"

**Supplementary**

**ImageJ Macro 1: WPB number, angle and feret diameter**

```
imageTitle=getTitle();//returns a string with the image title
run("Split Channels");
selectWindow("C1-"+imageTitle)
close(); selectWindow("C2-"+imageTitle);
setOption("ScaleConversions", true);
run("8-bit");
run("Subtract Background...", "rolling=3 sliding");
run("Auto Local Threshold", "method=Bernsen radius=15 parameter_1=0
parameter_2=0 white");
run("Analyze Particles...", "size=0.1-3.00 show=Masks clear add");
run("Set Measurements...", "area perimeter feret's redirect=None decimal=9");
roiManager("Measure");
saveAs("Results", "C:\\".txt");
selectWindow("C2-"+imageTitle);
close();
selectWindow("Mask of C2-"+imageTitle);
close();
close("Results");
close("ROI Manager");
```

**ImageJ Macro 2: VWF Strings**

```
imageTitle=getTitle();//returns a string with the image title run("Split
Channels");
selectWindow("C1-"+imageTitle)
close();
selectWindow("C2-"+imageTitle);
setOption("ScaleConversions", true);
run("8-bit");
run("Subtract Background...", "rolling=0.5 sliding");
run("Auto Local Threshold", "method=Phansalkar radius=15 parameter_1=0
parameter_2=0 white");
run("Analyze Particles...", "size=2-Infinity circularity=0.00-0.20 show=Masks
clear add exclude");
saveAs("Results", "C:\\ +".txt");
selectWindow("C2-"+imageTitle);
close();
selectWindow("Mask of C2-"+imageTitle);
close();
close("Results");
close("ROI Manager");
```
